# Differential characterization of physiological and biochemical responses during drought stress in finger millet varieties

**DOI:** 10.1101/603944

**Authors:** Asunta Mukami, Alex Ngetich, Cecilia Mweu, Richard O. Oduor, Mutemi Muthangya, Wilton Mwema Mbinda

**Author notes:** Corresponding authors: Asunta Mukami, Position, M.Sc. Student, E-mail addresses; Wilton Mbinda, Position, Lecturer, E-mail addresses.

## Abstract

Drought is the most perilous abiotic stress that affects finger millet growth and productivity worldwide. For the successful production of finger millet, selection of drought tolerant varieties is necessary and critical stages under drought stress, germination and early seedling growth, ought to be fully understood. This study investigated the physiological and biochemical responses of six finger millet varieties (GBK043137, GBK043128, GBK043124, GBK043122, GBK043094 and GBK043050) under mannitol-induced drought stress. Seeds were germinated on sterile soil and irrigated with various concentrations of mannitol (200, 400 and 600 mM) for two weeks. Comparative analysis in terms of relative water content (RWC), chlorophyll, proline, and malondialdehyde (MDA) contents were measured the physiological and biochemical characteristics of drought stress. The results showed that increased level of drought stress seriously decreased germination and early seedling growth of finger millet varieties. However, root growth was increased. In addition, exposition to drought stress triggered a significant decrease in relative water content and chlorophyll content reduction the biochemical parameters assay showed less reduction of relative water content. Furthermore, oxidative damage indicating parameters such as proline concentration and MDA content increased. Varieties GBK043137 and GBK043094 were less affected by drought as shown by significant change in the physiological parameters. Our findings reveal the difference and linkage between the physiological responses of finger millet to drought and are vital for breeding and selection of drought tolerant varieties of finger millet. Further investigations on genomic and molecular to deeply insight the detail mechanisms of drought tolerance in finger millet need to explored.

## Introduction

Drought stress, which mostly characterize arid and semi-arid regions of the world, is one of the most severe environmental stress which is responsible for poor agricultural productivity and yield decline (Zougmoré, 2018). The climate of most of sub-Saharan African is characterized by high temperature and low rainfall, during the vegetation seasons. (Rishmaw et al., 2016). Due to global climate change, it is predicted that drought episodes will increase in frequency, be longer and more severe, exacerbating its negative effects on crops and compromise food security particularly in developing countries. Over time, plants have evolved a range of drought tolerance adaptative mechanisms to counteract the detrimental effects of drought. When grown under desiccation stress, plants exhibit a sequence series of morphological, physiological, biochemical, cellular and molecular changes that severely compromise plant’s growth, development and productivity (Li and Liu, 2016). Plants under water deficit conditions decrease net photosynthesis and transpiration rates. These two physiological responses, which vary depending on the species, are often seen in regions with very high evaporative demand (Anjum et al., 2011). Protection systems against excess reducing power are therefore a vital approach for plants under desiccation stress (Chaves et al., 2009). Drought stress in plants is physiologically complex and it encompasses osmotic stress and specific ion toxicity (Todaka et al., 2015). Drought stress in plants is associated with nutritional imbalance, adjustment in metabolic fluxes, distortion and disorganization of cell and chloroplast membranes as well as reduction in division and expansion of cells and overproduction of reactive oxygen species (ROS) (Forni et al., 2017). Toxicity accruing from overproduction of ROS triggers cascades of oxidative reactions which consequently causes inactivation of enzymes and increase of lipid peroxidation, whose final product is malondialdehyde (MDA) and its quantification is used as a marker for oxidative damage (Moller et al., 2007). To abate the effects of oxidative stress, plants have evolved complex enzymatic and non-enzymatic systems. When exposed to water deficit stress conditions, many plant species enhance the activities of antioxidant enzyme which are associated with increased proline concentration (Ashraf and Foolad, 2007). Proline plays significant role in the osmoregulation, allowing cells to retain more water. Moreover, the amino acid also displays plant defense properties as a ROS scavenger (Szabados and Savouré, 2010) and as a regulator of the cellular redox status (Sharma et al., 2011). Proline accumulation in plants is therefore considered as a positive indicator for their tolerance to water stress (Verslues et al., 2014). Plants capability to retain water during desiccation is a vital strategy for plant tolerance to stress caused by water deficit stress. Accordingly, evaluation of relative water content change is the best representation and a fast approach to evaluating genetic differences in cellular hydration, plant water deficit and physiological water status after water deficit stress treatments (Sánchez-Rodríguez et al. 2010). The best effective approach of mitigating drought is the development of the tolerant crop varieties. Accordingly, it is important to identify the genetic resources with high tolerance and to understand the physiological and biochemical response mechanisms of drought tolerance in important cereal crops such as finger millet.

Finger millet, [*Eleusine coracana* (L.) Gaertn.], is a cereal crop which is cultivated semi-arid and arid regions of world under rain fed conditions (Thilakarathna & Raizada, 2015). The crop plays a significant role in food security in arid and semi-arid regions of sub-Saharan Africa and South Asia. Finger millet is therefore an ideal crop for reshaping food propensity of people due to its nutritional richness, high photosynthetic efficiency, and better tolerance to biotic and abiotic stresses than other crops (Kumar et al., 2016). As a member of the *Panicoideae* subfamily, finger millet acts as a model cereal crop for investigating basic biological processes. Although most of the finger millet varieties are considered to be drought tolerant when compared with other cereal crops, such as sorghum, maize, rice, barley and wheat, the crop is drought sensitive especially at early stages, especially if the first rains of the season are distant from each other. Genetic variations in response to drought stress have been showed in many plant relatives and among accessions within the same species. To our knowledge, there is no literature available which reports morphological, physiological and biochemical responses of finger millet to water deficit stress. We therefore investigated the physiological and biochemical mechanisms involved in six finger millet varieties, from distinct geographical zones in Kenya, under mannitol induced drought stress. Physiological and biochemical parameters were measured such as germination rate, shoot growth and root growth, relative water content (RWC), chlorophyll content, proline accumulation and lipid peroxidation.

## Materials and methods

### Plant material, growth conditions and germination assay

Finger millet varieties GBK043137, GBK043128, GBK043124, GBK043122, GBK043094 and GBK043050 obtained Kenya Agricultural and Livestock Research Organization, Gene Bank, Muguga, Kenya were used in this study. Seeds were sorted by handpicking of the healthy ones which were used for subsequent experiments. Selected seeds were washed with distilled water to remove dust and other particles. Germination assay was performed using 10 seeds of each variety. Seeds were planted in germination trays containing sterile soil to a depth of approximately 1 cm and irrigated with different concentrations of mannitol (200, 400 and 600 mM). The controls were irrigated with distilled water. Drought stress on was imposed on treatment groups by irrigating the seeds with various concentrations of mannitol at an interval of 3 days for two weeks. Observations on the rate of germination were scored on the 17^th^ day of treatment.

### Growth conditions drought treatment

Germinated finger millet seedlings were grown for 2 weeks under greenhouse conditions of 25±2 °C and 60-70% humidity, with a 16/8-h photoperiod provided by natural sunlight. The seedlings were subjected to osmotic stress by irrigating with mannitol (200, 400 and 600 mM) for 21 days at an interval of 3 days. Control plants were watered with distilled water. Shoot length and root length were measured after the experiment.

### Determination of relative water content

A leaf was excised from each plant on the 21^st^ day of water deficit stress. Immediately, the fresh weight (FW) of each leaflet was determined. Thereafter, the leaflet was immersed in double distilled water and incubated under normal room temperature for 4 hours. Afterwards, the leaflet was taken out, thoroughly wiped to remove the water on the blade surface and its weight measured to obtain turgid weight (TW). the leaflet was afterwards dried in an oven for 24 hours and its dry weight (DW) measured. The relative water content (RWC %) was calculated using the formula: RWC = [(FW -DW)/ (TW - DW)] × 100.

### Total chlorophyll content

Total chlorophyll (TC) content was determined using the method of described by Arnon (1949). Fresh leaves (0.2 g) of leaves plants were crushed in 80% acetone. The extract was centrifuged at 5000g for 3 minutes. The absorbance of the obtained supernatants was measured at 645 and 663 nm using 1240 UV-Vis Spectrophotometer (Shimadzu, Kyoto, Japan). The total chlorophyll content in each sample, expressed in mg/g fresh mass (FM) was calculated using the formula: TC = 20.2(A_645_) 8.02(A_663_) ×V/1000 × W where V corresponds to the volume of total extract per litre, W is the mass of the fresh material and A is the absorbance as 645 and 663 nm.

### Estimation of proline content

The amount of free proline in fresh plant leaves was determined as reported by Bates et al. (1973). Fresh leaf tissues (50 mg) from each variety and treatment was homogenized in 10 ml of 3% w/v sulphosalicylic acid and the homogenate filtrated. The resulting solution was mixed with acidic ninhydrin solution [40% (w/v) acidic ninhydrin (8.8 µM ninhydrin, 10.5 M glacial acetic acid, 2.4 M orthophosphoric acid), 40% (v/v) glacial acetic acid and 20% (v/v) of 3%(v/v) sulphosalicylic acid]. Thereafter, the reaction mixtures were put in a water bath at 100 °C for 60 minutes to develop colors. The reaction was terminated by incubating the mixtures in ice for 5 minutes. Toluene was added to separate chromophores. The optical density was measured at 520 nm using 1240 UV-Vis Spectrophotometer. Free proline content [µmol/g fresh weight (F. WT)] in leaf tissues was calculated from a standard curve made using 0-100 µg L-proline.

### Lipid peroxidation assay

Fresh upper second fully expended leaves (0.3 g) were harvested and homogenized in 0.1 % (w/v) trichloroacetic acid and the homogenates were centrifuged at 10,000 g for 15 minutes at 4 °C. The supernatant was mixed with 0.5 ml of 1.5 ml 0.5% thiobarbituric acid diluted in 20% trichloroacetic acid and the resulting mixture was heated to 95 °C for 25 minutes in water bath before incubating it on ice for 10 minutes. The absorbance was measured at 532 and 600 nm using UVmini-1240 UV-Vis Spectrophotometer with 1% thiobarbituric acid in 20% trichloroacetic acid as control. The amount of malondialdehyde (µmol/g FW) calculated as a measure of lipid peroxidation, was determined according to Heath and Packer, (1968).

### Statistics data analysis

The experiment was completely randomized block design with five replications of 10 plants. For germination and physiological assays, 10 seeds per replication were employed. Data collected were subjected to one-way analysis of variance (ANOVA) followed by a Fisher’s protected LSD test to compare the means. A confidence level was set at of 95% (p ≤ 0.05). All statistical procedures were performed using Minitab statistical computer softwarev.17.

## Results

### Effects of drought stress on seed germination

The results demonstrated that the gemination rate of the tested finger millet varieties was significantly influenced by seed variety and mannitol concentration (Table 1). Under untreated conditions, results showed that the highest gemination rate was recorded after 5 days in variety GBK043137 (83.75%) followed by varieties GBK043124, GBK043128, GBK043122 and GBK043050 whose gemination rates ranged from 65.0% to 72.5%, while GBK043094 recorded the lowest one at 51.25%. Seeds geminated in absence of stress treatment recorded superior gemination percentages. Imposition of increasing concentration of mannitol resulted to a decrease in germination percentage. The decline was significantly pronounced at 400 mM mannitol where 0% germination rate for varieties GBK043137, GBK043122, GBK043094 and GBK043050 were recorded while varieties GBK043124 and GBK043128 recorded 16.25% and 1.25% germination rates respectively (Table 1). Under moderate drought stress of 200 mM mannitol, variety GBK043137 recorded the highest germination rate of 41.25% compared to the other varieties whose germination rates ranged from 3.75% to 16.25% (Table 1). In severe osmotic pressure of 600 mM mannitol concentration, none of planted seeds were geminated. The average germination period under 0 mM mannitol concentration was 5.2 to 7.4 days for all varieties, while under 200 mM mannitol the germination interval was longer, ranging from 7.5 days to 13.6 days.

**Table 1.** Effects of mannitol on germination of six finger millet varieties.

| Variety | 0 mM | 200 mM | 400 mM | 600 mM |
| --- | --- | --- | --- | --- |
| GBK043137 | 83.75±4.7<sup>a</sup> | 41.25±8.75<sup>a</sup> | 0.00±0.00<sup>b</sup> | 0.00±0.00<sup>a</sup> |
| GBK043128 | 65.0±14.00<sup>ab</sup> | 3.75±2.39<sup>b</sup> | 1.25±1.25<sup>b</sup> | 0.00±0.00<sup>a</sup> |
| GBK043124 | 72.50±4.33<sup>ab</sup> | 16.25±3.75<sup>b</sup> | 16.25±8.26<sup>a</sup> | 0.00±0.00<sup>a</sup> |
| GBK043122 | 65.00±7.36<sup>ab</sup> | 3.75±2.39<sup>b</sup> | 0.00±0.00<sup>b</sup> | 0.00±0.00<sup>a</sup> |
| GBK043094 | 51.25±5.91<sup>b</sup> | 8.75±7.18<sup>b</sup> | 0.00±0.00<sup>b</sup> | 0.00±0.00<sup>a</sup> |
| GBK043050 | 66.25±9.66<sup>ab</sup> | 6.25±3.15<sup>b</sup> | 0.00±0.00<sup>b</sup> | 0.00±0.00<sup>a</sup> |

Means (±SE) followed by different alphabets in each column are significantly different (P≤0.05) using Fishers LSD

### Effects of drought stress on growth

The present study investigated the changes in the growth parameters (shoot and root growth) under mannitol induced drought conditions in all six finger millet varieties selected. The plant growth in the six varieties recorded remarkably higher responses in terms of shoot growth in absence of stress treatment compared to those exposed to mannitol induced drought stress (Fig. 1). The shoot length decreased progressively with increase in mannitol concentration (Table 2). Under mannitol stress conditions, higher growth responses were recorded at 200 mM mannitol, while the least responses were recorded at 600 mM mannitol (Table 2). Under stress conditions, variety GBK043128 recorded highest shoot length (3.00 cm) while the least response was observed in varieties GBK043137 and GBK043094 at 1.20 cm respectively (Table 2). Significance differences on the effect of mannitol on shoot length were only observed at 200 mM mannitol concentration. Higher mannitol concentrations did not record any significance differences among the varieties on shoot length (Table 2).

**Table 2.** Effect of mannitol on shoot length.

| Variety | 0 mM | 200 mM | 400 mM | 600 mM |
| --- | --- | --- | --- | --- |
| GBK043137 | 7.80±0.86<sup>a</sup> | 2.30±0.20<sup>b</sup> | 1.80±0.27<sup>a</sup> | 1.20±0.20<sup>a</sup> |
| GBK043128 | 7.60±1.33<sup>a</sup> | 3.00±0.27<sup>a</sup> | 2.20±0.26<sup>a</sup> | 1.30±0.20<sup>a</sup> |
| GBK043124 | 4.40±0.40<sup>b</sup> | 2.20±0.20<sup>b</sup> | 2.00±0.27<sup>a</sup> | 1.30±0.20<sup>a</sup> |
| GBK043122 | 4.00±0.45<sup>b</sup> | 2.40±0.29<sup>ab</sup> | 1.70±0.20<sup>a</sup> | 1.30±0.20<sup>a</sup> |
| GBK043094 | 3.00±0.00<sup>b</sup> | 2.40±0.19<sup>ab</sup> | 1.60±0.19<sup>a</sup> | 1.20±0.20<sup>a</sup> |
| GBK043050 | 3.70±0.62<sup>b</sup> | 2.10±0.10<sup>b</sup> | 1.60±0.19<sup>a</sup> | 1.30±0.20<sup>a</sup> |

Means (±SE) followed by different alphabets in each column are significantly different (P≤0.05) using Fishers LSD.

**Fig 1.**
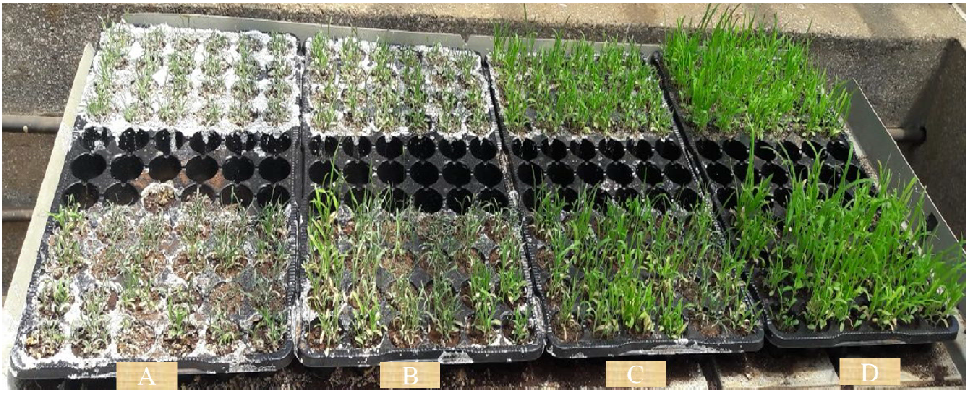
Effect of drought stress on growth of finger millet. Seedling growth on (A) 600 mM mannitol. (B 400 mM mannitol; (C) 200 mM mannitol; (D) 0 mM mannitol.

Contrary to shoot growth under mannitol osmotic stress conditions, the six finger millet varieties recorded an increase in root growth with increase in drought severity. The mannitol stressed plants recorded relatively higher responses when compared to control plants (Table 3). Variety GBK043094 recorded the highest root length under drought of 6.00 cm at 600 mM mannitol while GBK043050 and GBK043137 showed the least response with 2.30 cm and 2.60 cm respectively, at 200 mM mannitol treatment level (Table 3). The observed increase of root length across different drought stress levels was variety dependent.

**Table 3.** Effect of mannitol on root growth.

| Variety | 0 mM | 200 mM | 400 mM | 600 mM |
| --- | --- | --- | --- | --- |
| GBK043137 | 3.10±0.75<sup>a</sup> | 2.60±0.73<sup>b</sup> | 2.70±0.62<sup>a</sup> | 3.20±0.68<sup>c</sup> |
| GBK043128 | 3.20±0.37<sup>a</sup> | 4.30±0.49<sup>a</sup> | 4.60±0.93<sup>a</sup> | 5.00±0.45<sup>ab</sup> |
| GBK043124 | 2.60±0.40<sup>a</sup> | 3.20±0.37<sup>ab</sup> | 3.60±0.40<sup>a</sup> | 3.60±0.68<sup>bc</sup> |
| GBK043122 | 2.70±0.62<sup>a</sup> | 3.40±0.25<sup>ab</sup> | 3.60±0.68<sup>a</sup> | 5.00±0.45<sup>ab</sup> |
| GBK043094 | 2.0±0.57<sup>a</sup> | 3.50±0.78<sup>ab</sup> | 3.60±0.68<sup>a</sup> | 6.00±0.84<sup>a</sup> |
| GBK043050 | 2.00±0.61<sup>a</sup> | 2.30±0.30<sup>b</sup> | 2.90±0.56<sup>a</sup> | 3.90±0.25<sup>bc</sup> |

Means (±SE) followed by different alphabets in each column are significantly different (P ≤0.05) using Fishers LSD.

### Effects of drought stress on relative water content

Table 4 presents the RWC changes in finger millet leaves along with increase in water-deficit stress. Under irrigated conditions, all varieties maintained the highest RWC. Exposition of the plants to progressive mannitol concentrations simultaneously reduced RWC values of all varieties. The per cent reduction in RWC was the highest in GBK043122 which exhibited the lowest RWC value under water deficit stress at all the mannitol regimes. Variety GBK043128 sustained relatively high values of RWC and also showed lower percent reduction when compared to other varieties under water deficit stress. Plants under moderate water stress treatment of 200 mM mannitol displayed the highest diversity RWC values. The leaves exhibited wilting symptoms and leaf rolling at severe drought stress treatments.

**Table 4.** Effects of mannitol on relative water content (%)

| Variety | 0 mM | 200 mM | 400 mM | 600 mM |
| --- | --- | --- | --- | --- |
| GBK043137 | 85.56±4.12<sup>a</sup> | 68.60±5.27<sup>c</sup> | 64.96±4.62<sup>ab</sup> | 49.76±3.78<sup>ab</sup> |
| GBK043128 | 85.84±3.05<sup>a</sup> | 74.24±2.33<sup>b</sup> | 65.24±2.68<sup>ab</sup> | 54.76±4.23<sup>a</sup> |
| GBK043124 | 77.20±5.03<sup>ab</sup> | 67.14±3.02<sup>c</sup> | 60.78±4.88<sup>bc</sup> | 49.38±4.85<sup>b</sup> |
| GBK043122 | 74.16±2.94<sup>c</sup> | 66.92±3.05<sup>c</sup> | 57.98±4.06<sup>c</sup> | 40.18±1.96<sup>c</sup> |
| GBK042094 | 85.92±3.76<sup>a</sup> | 75.50±4.12<sup>b</sup> | 68.84±2.71<sup>a</sup> | 46.82±3.55<sup>b</sup> |
| GBK043050 | 81.94±7.91<sup>ab</sup> | 83.44±5.92<sup>a</sup> | 66.14±6.32<sup>ab</sup> | 48.74±5.28<sup>b</sup> |

Means (±SE) followed by different alphabets in each column are significantly different (P≤0.05) using Fishers LSD.

### Effects of drought stress on total chlorophyll content

Results from our study show an inverse relationship between mannitol induced drought stress responses and total chlorophyll content values for all finger millet varieties. Differences for chlorophyll content values were also observed among varieties. At the beginning of the experiment, total chlorophyll content across the varieties was similar ranging from 15.35 to 21.74 mg/g FW (Table 5). Imposition of moderate drought stress conditions of 200 mM mannitol caused a slight decrease of chlorophyll content ranging from 5.08% for GBK043094 to 14.2% for variety GBK043128. Significant decrease of ranging from 33.04 to 45.59% was observed at severe water stress conditions of 600 mM. Among the varieties exposed to severe water stress, varieties GBK043137 and GBK042094 retained relatively high chlorophyll content while drought-sensitive varieties GBK043050, GBK043128, GBK043122 and GBK043124 recorded a higher decline in chlorophyll reduction, ranging from 42.4% to 45.59% under mannitol induced drought stress (Table 5). The high drought-induced decrease of the total chlorophyll content signifies that drought stresses induced a high loss of photosynthetic reaction centers.

**Table 5.** Effects of mannitol on chlorophyll content (mg/g FW)

| Variety | 0 mM | 200 mM | 400 mM | 600 mM |
| --- | --- | --- | --- | --- |
| GBK043137 | 15.35±1.12<sup>b</sup> | 14.51±1.23<sup>c</sup> | 11.81±0.68<sup>b</sup> | 10.27±0.61<sup>abc</sup> |
| GBK043128 | 21.74±2.26<sup>a</sup> | 18.65±1.90<sup>a</sup> | 14.23±1.49<sup>a</sup> | 12.30±1.29<sup>a</sup> |
| GBK043124 | 17.33±1.47<sup>b</sup> | 15.16±1.78<sup>bc</sup> | 11.40±1.02<sup>b</sup> | 9.99±1.00<sup>b</sup>c |
| GBK043122 | 16.56±1.12<sup>b</sup> | 15.06±0.91<sup>bc</sup> | 10.96±1.03<sup>b</sup> | 9.76±1.58<sup>c</sup> |
| GBK042094 | 18.26±2.57<sup>b</sup> | 17.33±2.35<sup>ab</sup> | 14.32±2.15<sup>a</sup> | 12.14±1.78<sup>ab</sup> |
| GBK043050 | 16.78±0.07<sup>b</sup> | 14.86±0.06<sup>bc</sup> | 10.55±0.06<sup>b</sup> | 9.13±0.23<sup>c</sup> |

Means (±SE) followed by different alphabets in each column are significantly different (P≤0.05) using Fishers LSD.

### Effect of mannitol on proline content

The variations among the varieties in proline content under control conditions were significantly different and also did not follow any pattern (Table 6). In response to drought stress, all the varieties exhibited a steep increase in leaf proline content and the amount increased with the increased severity to the water stress. Variety GBK042094 had highest proline accumulation while GBK043128 had the least proline concentration in all mannitol treatments. Varietal differences in drought stress induced proline were clearly observed in finger millet, signifying a correlation between proline accumulation and differential mannitol induced water deficit stress tolerance response among the six finger millet varieties studied.

**Table 6.** Effects of mannitol on proline content (µmol/g FW)

| Variety | 0 mM | 200 mM | 400 mM | 600 mM |
| --- | --- | --- | --- | --- |
| GBK043137 | 1.76±0.09<sup>a</sup> | 2.12±0.19<sup>ab</sup> | 3.22±0.26<sup>a</sup> | 4.28±0.29<sup>a</sup> |
| GBK043128 | 1.76±0.27<sup>a</sup> | 1.90±0.16<sup>c</sup> | 2.76±0.21<sup>b</sup> | 3.76±0.18<sup>c</sup> |
| GBK043124 | 1.74±0.27<sup>a</sup> | 1.98±0.19<sup>abc</sup> | 2.84±0.17<sup>b</sup> | 3.50±0.14<sup>c</sup> |
| GBK043122 | 1.86±0.34<sup>a</sup> | 1.92±0.23<sup>bc</sup> | 2.86±0.21<sup>b</sup> | 3.80±0.17<sup>b</sup> |
| GBK042094 | 1.70±0.21<sup>a</sup> | 2.16±0.19<sup>a</sup> | 3.28±0.18<sup>a</sup> | 4.52±0.22<sup>a</sup> |
| GBK043050 | 1.74±0.27<sup>a</sup> | 1.98±0.15<sup>abc</sup> | 2.82±0.19<sup>b</sup> | 3.60±0.24<sup>bc</sup> |

Means (±SE) followed by different alphabets in each column are significantly different (P≤0.05) using Fishers LSD.

### MDA content

Lipid peroxidation was determined by measuring the accumulation of MDA, which is natural product of oxidation of polyunsaturated fatty acids present in the membrane caused by accumulation of peroxyl radicals (Kotchoni, et al. 2006). Our results revealed that the MDA levels in finger millet leaves was significantly influenced by severity of mannitol induced osmotic stress and variety. At the beginning of the experiment, no significant difference was registered in MDA values for all finger millet varieties (Table 7). The MDA content was lower in control plants ranging from 2.1 to 2.79 µmol/g FW compared to plants subjected to mannitol induced drought stress which ranged from 2.77 to 7.23 µmol/g FW. A progressive increase in the level of lipid peroxidation was observed with concomitant increase of mannitol concentration. The maximum MDA content under severe osmotic drought conditions (600 mM mannitol) was observed in GBK043128 followed by GBK043050 and GBK043122 varieties while varieties GBK042094 and GBK043137 had the least MDA accumulation at similar conditions (Table 7).

**Table 7.** Effects of mannitol on malondialdehyde content (µmol/g FW)

| Variety | 0 mM | 200 mM | 400 mM | 600 mM |
| --- | --- | --- | --- | --- |
| GBK043137 | 2.03±0.55<sup>c</sup> | 2.77±0.39<sup>c</sup> | 4.29±0.62<sup>d</sup> | 5.26±0.34<sup>c</sup> |
| GBK043128 | 2.27±0.46<sup>abc</sup> | 3.43±0.49<sup>b</sup> | 5.75±0.36<sup>a</sup> | 7.23±0.36<sup>a</sup> |
| GBK043124 | 2.58±0.33<sup>abc</sup> | 3.91±0.37<sup>ab</sup> | 5.00±0.45<sup>bc</sup> | 6.17±0.47<sup>b</sup> |
| GBK043122 | 2.66±0.38<sup>ab</sup> | 4.21±0.33<sup>a</sup> | 5.72±0.35<sup>a</sup> | 7.03±0.53<sup>a</sup> |
| GBK042094 | 2.79±0.63<sup>a</sup> | 3.74±0.67<sup>ab</sup> | 4.41±0.77<sup>cd</sup> | 5.39±0.51<sup>c</sup> |
| GBK043050 | 2.10±0.15<sup>bc</sup> | 3.63±0.27<sup>b</sup> | 5.67±0.60<sup>ab</sup> | 7.62±0.97<sup>a</sup> |

Means (±SE) followed by different alphabets in each column are significantly different (P≤0.05) using Fishers LSD.

## Discussion

Drought stress induces different physiological, genetic and metabolic responses among several species of plant and varieties. These responses are also influenced by edaphic, climatic and agronomic factors (Caliz et al., 2015). Vulnerability of plants to drought stress differentially varies in depending on stress severity, interactions among stressors, plant species and stages of their development (Demirevska et al., 2009). This natural allelic difference may provide valuable information into the mechanisms which underline the differential responses to agriculturally important traits and search of the crops that can survive such harsh environments may assist to ensure stable and sustainable food production (Budak et al., 2013). As a dry-land crop, finger millet growth and productivity is highly affected by drought stress which is projected to increase in severity and frequency with current adverse climate change era. In order to overcome this, there is need to develop new finger millet varieties with strong drought tolerance traits as an effective way to achieve high and stable yields. For this to be successful, precise identification of stress tolerance of finger millet varieties forms the basis of developing resistant finger millet varieties. Therefore, dissecting the natural differences of finger millet varieties could be viable to explore the complex mechanisms of its response to various stresses. This study was done to investigate the differential responses of finger millet to seed germination, growth, physiological and biochemical responses after exposure to different concentrations of mannitol, which causes osmotic stress and is commonly used as a drought simulator (Ullah et al., 2014; Kaya et al., 2013; Karakas et al., 1997).

In plants life cycle, seed germination is the most critical and sensitive stage. The process of seed gemination is constrained or even completely prevented by drought stress (Hubbard et al. 2012). Germination potential is therefore an ideal index which is used to assess the seed germination rate and germination uniformity. The germination rate under simulated drought stress showed the tolerance, though the responses were variety dependent. In absence of stress treatment, the six finger millet varieties recorded better germination percentages. However, the rate declined with increase in mannitol concentration treatment. Similar results have been reported in other plant species such as maize (Liu et al. (2015), wheat (Yang et al., 2016) and sunflower (Ahmad et al., 2009). Seed germination process is divided into three successive stages: inhibition, metabolism that leads initiation of radicle growth, and radicle growth which primes radicle emergence. A threshold level of hydration is essential for the ensuing radicle elongation (Ramagopal, 1990). In normal seed germination process, a threshold of the embryo hydration level needs to be attained, which is a critical precondition for the successive initiation of cell elongation and radicle development (Hegarty, 1978). In our study, the presence of mannitol could have severely reduced internal osmotic potential of the germinating seeds, therefore permitting the water uptake which subsequently leads to germination initiation processes.

Plants capability to retain high water status during desiccation stress is a vital strategy for plant tolerance to drought stress. Accordingly, evaluation of relative water content change is the best representation and a fast approach to evaluating genetic differences in cellular hydration, plant water deficit and physiological water status after water deficit stress treatments (Sánchez-Rodríguez et al. 2010). Normally, high relative water content values are treated as index of drought stress tolerance, as demonstrated by Pandey et al. (2015) on rice genotypes tolerant or sensitive to drought. The differences in relative water content in all varieties observed in our study could be correlated with their different ability of water absorption from soil. The decline in relative water content recorded was a main factor that caused decreased growth responding to osmotic stress in the finger millet plants. Under desiccation stress, sensitive finger millet varieties were more affected by the decrease in relative water content than tolerant varieties. This suggested that the six finger millet varieties had different sensitivity when subjected mannitol induced drought. The enhanced water retention capacity observed in some of finger millet even when challenged by drought could play a vital role in for plant survival under drought conditions water deficit.

Plants chlorophyll content heavily depends on the species physiological responses and their ability to resist environmental stresses (Anjum et al., 2011). Evaluation of leaf chlorophyll concentration is one of the most effective diagnostic tool for studies of drought tolerance identification, genotypic variation, altitudinal variation and has been employed in many crops including cereals such as sorghum (Qadir et al., 2015) and foxtail millet (Wang et al., 2016). Plants can overcome this assault by increasing the accumulation of chlorophyll which protects the plants by getting rid of excessive energy by thermal dissipation (Reddy et al., 2004). Consequently, decline of chlorophyll concentration in response to drought stress is a common phenomenon, occasioned by disordering chlorophyll synthesis and resulting to plant chlorosis. Additionally, when plants are subjected to environmental stresses, leaf chloroplasts are injured which leads to disrupted photosynthesis. At higher mannitol concentrations above 200 mM, chlorosis was observed in all the varieties, and the leaves turned into pale yellow which lead to plant death.

Proline plays significant role in the osmoregulation, allowing cells to retain more water. Moreover, the amino acid also displays plant defense properties as a ROS scavenger (Szabados and Savouré, 2010) and as a regulator of the cellular redox status (Sharma et al., 2011). Proline accumulation has therefore a positive connection with their tolerance to various environmental stresses (Szabados and Savouré, 2010). In our study, the mannitol stressed plants showed significantly higher proline concentration was than control plants, especially in GBK042094. Our results revealed that free proline accumulation in the leaf tissues of drought susceptible finger millet varieties was significantly lower than the tolerant ones. These findings are corroborated by the data reported in previous research work which indicate that total free proline in the leaves are higher in water deficit tolerant than in drought susceptible lines of maize (Efeoğlu et al., 2009), sweetpotato (Mbinda et al., 2018), and rice (Pandey et al., 2015). The responses across the plant lines were concomitantly increased with progressive increment of mannitol dosage. Our results suggest that higher proline content in drought tolerant finger millet lines could be due to altered expression of drought responsive genes which potentially improve the hydration status of the plants. Our results also reinforce a close association between increased proline concentration and plant relative water content in drought tolerance mechanisms.

It is vital for antioxidative systems of plants to scavenge excess ROS in order to maintain a balanced equilibrium of cellular reactions when they challenged stress conditions (van Breusegem et al., 2018). The toxicity of ROS is due to their reactions with numerous cell components, which cause lipid peroxidation among other cascades of oxidative reactions (Wang et al., 2012). Cellular lipid peroxidation damages the plasma membrane, leading to leakage of contents, swift desiccation and cellular death (Demidchik, 2015). The final product of lipid peroxidation, is malondialdehyde and this solute is one of the best physiological biomarkers of drought tolerance in plants (Anjum et al., 2011). In this work, we found varieties and GBK043137 and GBK043094 having the least amounts of MDA when challenged by drought stress (Table 7). Low MDA levels has been correlated with desiccation stress tolerance and the ensuing lipid peroxidation could induce the activity of antioxidant enzymes (Wang et al., 2012). Accumulation of MDA when challenged by environmental stresses has also been found to be a good drought tolerance index in other plant species pitanga (Toscano et al., 2016), melon (Sarabi, et al., 2017), desi chickpea (Farooq et al., 2018) and wheat (Mickky and Aldesuquy, 2016). From all the physiological responses examined, it evident that of finger millet responses to drought stress largely depends on the genotype/cultivars used the length and severity of water deficit stress and the stage of development of the plant.

## Conclusion

In conclusion, our study provided a broad analysis of the physiological features of several finger millet plants to drought stress. The results reported here demonstrate the impact of drought stress on the analysed parameters with a wide range of variability among the studied varieties. Finger millet varieties GBK042094 and GBK043137 could tolerate water deficit better than four the other varieties, as indicated by significant decreases in germination rate, shoot length, root growth, relative water content, leaf total chlorophyll content, proline accumulation and lipid peroxidation. We deduced that these varieties are promising resources with considerable level of tolerance to drought stress and they can be used for further evaluations and breeding programs. Further investigations on genomic and molecular to deeply insight the detail mechanisms of drought tolerance in finger millet need to explored.

## Contributions

AM and AN carried all the experiments. AM helped with draft the manuscript. CM, RO, MM and WM supervised the study, contributed in statistical analysis and writing the manuscript, WM conceived the idea, obtained of funding, contributed with experimental design, coordination and manuscript writing. All authors agreed on the final appearance of the manuscript after careful review.

## Acknowledgement

This work was supported financially by a grant from The World Academy of Sciences (Ref. No. 17-357 RG/BIO/AF/AC_I – FR3240297745) through the generous contribution of the Swedish International Development Cooperation Agency. The authors gratefully acknowledge Kenya Agricultural and Livestock Research Organization, Gene Bank, for providing the finger millet seeds used. The authors also thank the Department of Biochemistry and Biotechnology, Kenyatta University for providing facilities for the research.

## References

Ahmad, S., Ahmad, R., Ashraf, M. Y., Ashraf, M., & Waraich, E. A. (2009). Sunflower (Helianthus annuus L.) response to drought stress at germination and seedling growth stages. Pak. J. Bot, 41(2), 647–654.

Anjum, S. A., Xie, X. Y., Wang, L. C., Saleem, M. F., Man, C., & Lei, W. (2011). Morphological, physiological and biochemical responses of plants to drought stress. African Journal of Agricultural Research, 6(9), 2026–2032.

Arnon, D. I. (1949). Copper enzymes in isolated chloroplasts. Polyphenoloxidase in *Beta vulgaris*. Plant physiology, 24(1), 1.

Arnon, D. I. (1949). Copper enzymes in isolated chloroplasts. Polyphenoloxidase in Beta vulgaris. Plant physiology, 24(1), 1.

Ashraf, M. F. M. R., & Foolad, M. (2007). Roles of glycine betaine and proline in improving plant abiotic stress resistance. Environmental and experimental botany, 59(2), 206–216.

Bates, L. S., Waldren, R. P., & Teare, I. D. (1973). Rapid determination of free proline for water-stress studies. Plant and soil, 39(1), 205–207.

Budak, H., Kantar, M., & Kurtoglu, K. (2013). Drought tolerance in modern and wild wheat. The Scientific World Journal, 2013.

Caliz, J., Montes-Borrego, M., Triadó-Margarit, X., Metsis, M., Landa, B. B., & Casamayor, E. O. (2015). Influence of edaphic, climatic, and agronomic factors on the composition and abundance of nitrifying microorganisms in the rhizosphere of commercial olive crops. PloS one, 10(5), e0125787.

Chaves, M. M., & Oliveira, M. M. (2004). Mechanisms underlying plant resilience to water deficits: prospects for water-saving agriculture. Journal of experimental botany, 55(407), 2365–2384.

Demidchik, V. (2015). Mechanisms of oxidative stress in plants: from classical chemistry to cell biology. Environmental and experimental botany, 109, 212–228.

Demirevska, K., Zasheva, D., Dimitrov, R., Simova-Stoilova, L., Stamenova, M., & Feller, U. (2009). Drought stress effects on Rubisco in wheat: changes in the Rubisco large subunit. Acta Physiologiae Plantarum, 31(6), 1129.

Efeoğlu, B., Ekmekci, Y., & Cicek, N. (2009). Physiological responses of three maize cultivars to drought stress and recovery. South African Journal of Botany, 75(1), 34–42.

Ekanayake, I. J., O’toole, J. C., Garrity, D. P., & Masajo, T. M. (1985). Inheritance of root characters and their relations to drought resistance in rice 1. Crop Science, 25(6), 927–933.

Ellouzi, H., Hamed, K. B., Cela, J., Munné□Bosch, S., & Abdelly, C. (2011). Early effects of salt stress on the physiological and oxidative status of *Cakile maritima* (halophyte) and *Arabidopsis thaliana* (glycophyte). Physiologia Plantarum, 142(2), 128–143.

Farooq, M., Ullah, A., Lee, D. J., Alghamdi, S. S., & Siddique, K. H. (2018). Desi chickpea genotypes tolerate drought stress better than kabuli types by modulating germination metabolism, trehalose accumulation, and carbon assimilation. Plant Physiology and Biochemistry, 126, 47–54.

Forni, C., Duca, D., & Glick, B. R. (2017). Mechanisms of plant response to salt and drought stress and their alteration by rhizobacteria. Plant and Soil, 410(1-2), 335–356.

Heath, R. L., & Packer, L. (1968). Photoperoxidation in isolated chloroplasts: I. Kinetics and stoichiometry of fatty acid peroxidation. Archives of biochemistry and biophysics, 125(1), 189–198.

Hegarty, T. W. (1978). The physiology of seed hydration and dehydration, and the relation between water stress and the control of germination: a review. Plant, Cell & Environment, 1(2), 101–119.

Hubbard, M., Germida, J., & Vujanovic, V. (2012). Fungal endophytes improve wheat seed germination under heat and drought stress. Botany, 90(2), 137–149.

Karakas, B., Ozias□Akins, P., Stushnoff, C., Suefferheld, M., & Rieger, M. (1997). Salinity and drought tolerance of mannitol□accumulating transgenic tobacco. Plant, Cell & Environment, 20(5), 609–616.

Kaya, C., Sonmez, O., Aydemir, S., Ashraf, M., & Dikilitas, M. (2013). Exogenous application of mannitol and thiourea regulates plant growth and oxidative stress responses in salt-stressed maize (*Zea mays* L.). Journal of plant interactions, 8(3), 234–241.

Kotchoni, S. O., Kuhns, C., Ditzer, A., Kirch, H. H., & Bartels, D. (2006). Over□expression of different aldehyde dehydrogenase genes in *Arabidopsis thaliana* confers tolerance to abiotic stress and protects plants against lipid peroxidation and oxidative stress. Plant, cell & environment, 29(6), 1033–1048.

Kumar, A., Metwal, M., Kaur, S., Gupta, A.K., Puranik, S., Singh, S., Singh, M., Gupta, S., Babu, B.K., Sood, S. & Yadav, R. (2016). Nutraceutical value of finger millet [*Eleusine coracana* (L.) Gaertn.], and their improvement using omics approaches. Frontiers in plant science, 7, p.934.

Li, X., & Liu, F. (2016). Drought stress memory and drought stress tolerance in plants: biochemical and molecular basis. In Drought Stress Tolerance in Plants, Vol 1 (pp. 17–44). Springer, Cham.

Liu, M., Li, M., Liu, K., & Sui, N. (2015). Effects of drought stress on seed germination and seedling growth of different maize varieties. Journal of Agricultural Science, 7(5), 231.

Mbinda, W., Ombori, O., Dixelius, C., & Oduor, R. (2018). Xerophyta viscosa Aldose Reductase, XvAld1, Enhances Drought Tolerance in Transgenic Sweetpotato. Molecular biotechnology, 60(3), 203–214.

Mickky, B. M., & Aldesuquy, H. S. (2017). Impact of osmotic stress on seedling growth observations, membrane characteristics and antioxidant defense system of different wheat genotypes. Egyptian Journal of Basic and Applied Sciences, 4(1), 47–54.

Møller, I. M., Jensen, P. E., & Hansson, A. (2007). Oxidative modifications to cellular components in plants. Annu. Rev. Plant Biol., 58, 459–481.

Pandey, V., & Shukla, A. (2015). Acclimation and tolerance strategies of rice under drought stress. Rice Science, 22(4), 147–161.

Pandey, V., Ansari, M. W., Tula, S., Yadav, S., Sahoo, R. K., Shukla, N., … & Kumar, A. (2016). Dose-dependent response of *Trichoderma harzianum* in improving drought tolerance in rice genotypes. Planta, 243(5), 1251–1264.

Passioura, J. B. (1983). Roots and drought resistance. In Developments in Agricultural and Managed Forest Ecology (Vol. 12, pp. 265–280). Elsevier.

Qadir, M., Quillérou, E., Nangia, V., Murtaza, G., Singh, M., Thomas, R.J., Drechsel, P. and Noble, A.D. (2014). November. Economics of salt□induced land degradation and restoration. In Natural Resources Forum (Vol. 38, No. 4, pp. 282–295.

Radhouane, L. (2007). Response of Tunisian autochthonous pearl millet (*Pennisetum glaucum* (L.) R. Br.) to drought stress induced by polyethylene glycol (PEG) 6000. African journal of biotechnology, 6(9),1102–1105.

Ramagopal, S. (1990). Inhibition of seed germination by salt and its subsequent effect on embryonic protein synthesis in barley. Journal of Plant Physiology, 136(5), 621–625.

Reddy, A. R., Chaitanya, K. V., & Vivekanandan, M. (2004). Drought-induced responses of photosynthesis and antioxidant metabolism in higher plants. Journal of plant physiology, 161(11), 1189–1202.

Rishmawi, K., Prince, S., & Xue, Y. (2016). Vegetation responses to climate variability in the northern arid to sub-humid zones of Sub-Saharan Africa. Remote Sensing, 8(11), 910.

Sánchez-Rodríguez, E., Rubio-Wilhelmi, M., Cervilla, L. M., Blasco, B., Rios, J. J., Rosales, M. A., & Ruiz, J. M. (2010). Genotypic differences in some physiological parameters symptomatic for oxidative stress under moderate drought in tomato plants. Plant science, 178(1), 30–40.

Sarabi, B., Bolandnazar, S., Ghaderi, N., & Ghashghaie, J. (2017). Genotypic differences in physiological and biochemical responses to salinity stress in melon (*Cucumis melo* L.) plants: Prospects for selection of salt tolerant landraces. Plant Physiology and Biochemistry, 119, 294–311.

Sharma, S., Villamor, J. G. C., & Verslues, P. E. (2011). Essential role of tissue specific proline synthesis and catabolism in growth and redox balance at low water potential. Plant physiology, pp-111.

Szabados, L., & Savoure, A. (2010). Proline: a multifunctional amino acid. Trends in plant science, 15(2), 89–97.

Thilakarathna, M. S., & Raizada, M. N. (2015). A review of nutrient management studies involving finger millet in the semi-arid tropics of Asia and Africa. Agronomy, 5(3), 262–290.

Todaka, D., Shinozaki, K., & Yamaguchi-Shinozaki, K. (2015). Recent advances in the dissection of drought-stress regulatory networks and strategies for development of drought-tolerant transgenic rice plants. Frontiers in plant science, 6, 84.

Toscano, S., Farieri, E., Ferrante, A., & Romano, D. (2016). Physiological and biochemical responses in two ornamental shrubs to drought stress. Frontiers in plant science, 7, 645.

Ullah, I., Akhtar, N., Mehmood, N., Shah, I. A., & Noor, M. (2014). Effect of mannitol induced drought stress on seedling traits and protein profile of two wheat cultivars. J. Animal Plant Sci, 24, 1246–1251.

Van Breusegem, F., Foyer, C., & Mann, G. (2018). Reactive oxygen species are crucial “pro-life “survival signals in plants. Free Radical Biology and Medicine 122: 1–3.

Wang, Y., Li, L., Tang, S., Liu, J., Zhang, H., Zhi, H., Jia, G & Diao, X. (2016). Combined small RNA and degradome sequencing to identify miRNAs and their targets in response to drought in foxtail millet. BMC genetics, 17(1), 57.

Yang, H., Feng, J., Zhai, S., Dai, Y., Xu, M., Wu, J., Shen, M., Bian, X., Koide, R.T. & Liu, J. (2016). Long-term ditch-buried straw return alters soil water potential, temperature, and microbial communities in a rice-wheat rotation system. Soil and Tillage Research, 163, 21–31.

Zougmoré, R. (2018). Promoting climate-smart agriculture through water and nutrient interactions options in semi-arid West Africa: A review of evidence and empirical analysis. In improving the profitability, sustainability and efficiency of nutrients through site specific fertilizer recommendations in West Africa agro-ecosystems (pp. 249–263). Springer, Cham.

